# Cannabis use, depression and self-harm: phenotypic and genetic relationships

**DOI:** 10.1101/549899

**Authors:** K Hodgson, JRI Coleman, SP Hagenaars, KL Purves, K Glanville, SW Choi, P O’Reilly, G Breen, Major Depressive Disorder Working Group of the Psychiatric Genomics Consortium, CM Lewis

## Abstract

**Background and Aims:** The use of cannabis has previously been linked to both depression and self-harm, however the role of genetics in this relationship are unclear. We aimed to examine the phenotypic and genetic relationships between these traits.

**Design:** Genetic and cross-sectional phenotypic data collected through UK Biobank, together with consortia genome-wide association study summary statistics. These data were used to assess the phenotypic and genetic relationship between cannabis use, depression and self harm.

**Setting:** UK, with additional international consortia data

**Participants:** N=126,291 British adults aged between 40 and 70 years, recruited into UK Biobank

**Measurements:** Genome-wide genetic data, phenotypic data on lifetime history of cannabis use, depression and self-harm.

**Findings:** In UK Biobank, cannabis use is associated with increased likelihood of depression (OR=1.64, 95% CI=1.59-1.70, p=1.19×10^−213^) and self-harm (OR=2.85, 95% CI=2.69-3.01, p=3.46×10^−304^). The strength of this phenotypic association is stronger when more severe trait definitions of cannabis use and depression are considered. Additionally, significant genetic correlations are seen between cannabis use and depression using consortia summary statistics (rg=0.289, SE=0.036, p=1.45×10^−15^). Polygenic risk scores for cannabis use and depression both explain a small but significant proportion of variance in cannabis use, depression and self harm within a UK Biobank target sample. However, two-sample Mendelian randomisation analyses were not significant.

**Conclusions:** Cannabis use is both phenotypically and genetically associated with depression and self harm. Future work dissecting the causal mechanism linking these traits may have implications for cannabis users.

## Introduction

Cannabis is one of the most commonly used psychoactive drugs^1^ and depression is the second leading cause of disability^2^ worldwide. Given the known comorbidity between these traits^3–7^, understanding the nature of their relationship is highly relevant from a public health perspective. This is particularly true in the context of changing legislation and public perceptions of the risks of using cannabis^8^.

Several theories regarding the mechanisms for associating cannabis use and depression have been proposed. One trait might causally lead to the other, or both traits could result from common factors (either genetic or environmental). There is some longitudinal evidence that cannabis use is associated with increased likelihood of later depression, however there are complexities in accounting for potential confounding factors^5,9^. Indeed, when all covariates are accounted for, cannabis use may follow rather than precede depression^10^. Thus, longitudinal attempts to disentangle causality remain inconclusive. However, this work highlights the potential impact of common risk factors that increase the likelihood of both using cannabis and suffering from depression.

Genetics is a useful tool by which to understand these common risk factors. Both cannabis use^11^ and depression^12^ are known to be heritable and substantial increases in sample sizes mean that there has been progress in understanding the genetic basis of these phenotypes individually. For cannabis use, the largest genome-wide association (GWA) study to date reported 8 significant loci^13^ and for depression, 44 genome-wide significant loci were identified by the Psychiatric Genomics Consortium Major Depressive Disorder (PGC-MDD) working group and 23andMe^14^. These developments mean efforts to understand the interplay between comorbidity traits from a genetic perspective are now much more powerful.

But there is mixed evidence for a shared genetic aetiology between cannabis use and depression. A significant genetic correlation between cannabis use and depression was observed in an extended pedigree sample^15^ but with a discordant twin design, genetic factors played only a small role in the link between early or frequent cannabis use and depression or suicidal ideation^16^. In unrelated samples, polygenic risk for depression was associated with non-problematic cannabis use^17^, however using genome-wide summary statistics to consider the genetic correlations between cannabis use and a range of genetic traits, the estimated correlation with depression was modest but not significant^13^. In the latter two analyses, the underpowered (and since superseded) PGC-MDD 2013 depression sample^18^ was used, which is likely to have impacted upon results. The recent availability of larger datasets enables us to examine the genetic relationship between cannabis use and depression with substantially more statistical power.

Cannabis use has also been associated with increased rates of self-harm; both for non-suicidal^19–21^ and suicidal^16,22,23^ behaviours. This phenotypic association may be stronger than that observed between cannabis and depression but whilst some studies report evidence of a shared genetic aetiology^23,24^ and others find that genetic factors are less important^16,25^. Therefore, additional work exploring how cannabis use related to self-harm is also needed.

Here we have chosen to focus on the use, rather than abuse of cannabis. Problematic cannabis use is more heritable^11^ and also potentially more strongly associated with depression^5^. But cannabis is not highly addictive^26^; estimates suggest only 10% of those who use cannabis become dependent^27^ meaning that cannabis use disorders are relatively rare. Further, they are an undesirable outcome in themselves. In the context of changing perceptions of the risks associated with cannabis use, understanding if and how mental health traits relate to non-disordered use of cannabis is arguably a more useful question from a public health perspective.

Here we use data from UK Biobank^28,29^ in addition to summary statistics available from genome-wide analyses by the PGC-MDD working group^14^ and the International Cannabis Consortium^30^ to explore the relationship between cannabis use, depression and self-harm. Many GWA studies of cannabis use consider all individuals who have ever tried cannabis use. Nevertheless, cross-sectional evidence suggests that more frequent (non-disordered) cannabis use is more strongly associated with depression^7^. Using the phenotypic detail available in UK Biobank, we investigate the risks associated with initial and continued cannabis use, as well as single episode and recurrent depression. We assess phenotypic associations and use genetic approaches^31–33^ to consider overlapping genetic influences on these traits. Finally, we use Mendelian randomisation (MR) techniques to examine the direction of causality between cannabis use and depression.

## Methods

### UK Biobank Sample

UK Biobank (http://www.ukbiobank.ac.uk) includes measures of many health-related phenotypes for approximately 500,000 British adults aged between 40 and 70 years^28^. UK Biobank received ethical approval from the Research Ethics Committee (11/NW/0382). The work presented here uses data from a subset of 157,366 participants who completed an online follow-up mental health questionnaire (category 136 on http://biobank.ctsu.ox.ac.uk), which included items on cannabis use, depression and self-harm^34^.

### Phenotypic Measurements

#### Cannabis

Participants were asked about their lifetime cannabis use (data field 20453): “Have you taken cannabis (marijuana, grass, hash, ganja, blow, draw, skunk, weed, spliff, dope), even if it was a long time ago?”. Those who responded “No” were classified as controls and those endorsing “Yes” options were classified as cannabis users. We separated these users into two groups: those reporting initial cannabis use (“Yes, 1-2 times”, “Yes, 3-10 times”) and continued cannabis use (“Yes, 11-100 times”, “Yes, more than 100 times”).

#### Depression

Using the work from Davis et al.^34^ as a framework, depression was defined using lifetime criteria based on questions derived from the Composite International Diagnostic Interview (CIDI). Exclusions were made if depression cases reported previous diagnoses of schizophrenia, other psychoses or bipolar disorder. However we did not exclude controls based on current prescriptions for antidepressant medications, given the use of these drugs for diagnoses other that depression. The reported number of depressed periods was used to classify individuals with single episode or recurrent depression.

#### Self harm

Participants were asked “Have you deliberately harmed yourself, whether or not you meant to end your life?” (data field 20480).

#### Other phenotypes

To consider potential confounding due to the use of substances other than cannabis, we used the “Ever smoked” item (derived data field 20160) as well as the mental health questionnaire item on “Frequency of drinking alcohol” (data field 20414).

### Genetic data

Genome-wide data is available for the UK Biobank cohort^29^. Genotyping was performed using two highly-overlapping custom genotyping arrays, covering ∼800,000 markers. This data underwent centralised quality control to remove genotyping errors before being imputed in a two-stage imputation to the Haplotype Reference Consortium^35^ and UK10K^36^ reference panels. Additional genotypic and sample quality controls thresholds were then applied and the sample was limited to individuals of White European ancestry (Supplementary Materials Section 1.2).

This gave a sample of N=126,291 individuals with both genetic and mental health questionnaire data available.

### Statistical methods

#### Univariate analyses within UK Biobank

##### Phenotypic analyses

For each of the seven phenotypes of interest (cannabis use, initial cannabis use, continued cannabis use, depression, single episode depression, recurrent depression and self-harm), chi-squared models were used to compare differences between cases and controls in terms of sex, smoking and frequency of alcohol drinking.

### Genome-wide association analyses

Using BGENIE v1.2^29^, genome-wide association (GWA) analyses were completed for each of the seven phenotypes, after residualising for assessment centre, batch and the first 6 principal components, as calculated in the European-only subset of the data using flashpca2^37^. Manhattan and QQ plots were made using the qqman^38^ R package. All positions are given in GRCh37 coordinates and results were clumped in plink 1.90^39^ (www.coggenomics.org/plink/1.9/) using clumping thresholds as in the PGC-MDD analysis^14^.

### Heritability of phenotypes

Using the generated GWA results and LD-score regression^31,32^, the SNP-heritability of each of the seven traits was calculated. For conversion to the liability scale, sample prevalence was defined as [Ncases/(Ncases + Ncontrols)], whilst population prevalence was defined as [Ncases]/[Ntotal completing MHQ]. To consider if the UK Biobank may have over-or under-sampled cases, we also calculated the heritability using population prevalence estimates plus or minus 20% of the cases observed.

### Bivariate analyses

#### Phenotypic relationships within UK Biobank

To explore the association between cannabis, depression and self-harm phenotypes, odds ratios were calculated using logistic regression models. To assess the impact of differences in sex, smoking and frequency of alcohol drinking, stratified analyses were perfomed.

### Genetic Correlations

The genetic relationships between phenotypes were examined using bivariate LD-score regression^31,32^.

#### Maximum available sample size

To consider the relationship between cannabis use and depression in the largest available sample, we used consortium-based summary statistics. For cannabis use, the International Cannabis Consortium (ICC, N=32,330^30^) is the largest publically available dataset. A meta-analysis combining the ICC with UK Biobank (UKB) was performed, total sample size N=158,621. Summary statistics from the ICC and UKB samples were cleaned using the EasyQC software^40^ and then meta-analysed using a p-value-based approach, with genomic control in METAL^41^. For this ICC+UKB combined cohort, the sample prevalence of cannabis use was 28%. For depression, the summary statistics from the PGC-MDD GWA were used, including both the UK Biobank and 23andMe cohorts (N=480,359, sample prevalence 28%^14^).

#### Detailed phenotypes within UK Biobank

To consider whether genetic correlations vary according the severity of the phenotype considered, bivariate LD-score analyses within UK Biobank were also performed, considering each combination of the seven traits of interest.

### Polygenic risk scoring

Consortium-based GWA summary statistics were also used to build polygenic risk scores for prediction into UK Biobank. Polygenic risk scoring can be biased by overlap between discovery and target samples. Therefore, when building a depression polygenic risk score (PRS_DEP_), UK Biobank was excluded from the PGC-MDD and 23andMe summary statistics in the discovery dataset (N=450,619). To maximise the discovery sample size available when building a polygenic risk score for cannabis use (PRS_CANN_), but retain an independent target sample for prediction, UK Biobank was split equally into two using the caret R package^42^ to create a discovery and a target subsample. The ICC data was then meta-analysed (using the same methods as described above) with the discovery 50% of UK Biobank to give a total discovery dataset for cannabis use of N=95,384 (ICC+50%UKB).

The target 50% of UK Biobank (N=63,054) was then used as the target sample for both polygenic risk scoring analyses, to facilitate comparison between phenotypes.

PRS scores were built using PRSice2 (https://choishingwan.github.io/PRSice/)^33,43^. For computational reasons, in the UK Biobank target dataset only directly genotyped variants were used. Full details are in Supplementary Materials Section 4, but briefly, 11 p-value thresholds were used when building the polygenic risk scores and a glm logistic model was used to test these. All R^2^ values shown have been converted to the liability scale^44^.

### Mendelian randomisation

To explore the direction of causality between cannabis use and depression, we used two-sample MR, using published genetic variants for cannabis use and depression as instrumental variables. MR can be biased by sample overlap. To obtain the largest sample size with available GWA summary statistics, whilst minimising sample overlap, we used the PGC-MDD with 23andMe (N=480,359) and ICC (N=32,330) datasets. Both of these datasets contain samples from the Netherlands Twin Register/NESDA^45,46^ and Queensland Institute of Medical Research Berghofer cohorts^18,47^. This gives a maximum possible overlap of 1.4% (N=6,719). Whilst results should be considered in the context of this overlap, Burgess et al.^48^ conclude that the bias from an overlap of this magnitude is minimal.

We used the R package TwoSampleMR^49^ to test casuality in both directions, using both inverse-variance weighted (IVW) and MR-Egger models (to consider evidence for pleiotropy). This package includes harmonisation steps to ensure allelic alignment between exposure and outcome datasets.

#### Testing the causal effect of depression on cannabis use

For the PGC-MDD and 23andMe dataset (SNP-h^2^=8.7%), 44 genome-wide significant independent loci have been previously identified^14^. The logOR and SE for the 39 of these SNPs were available in the ICC dataset.

#### Testing the causal effect of cannabis use on depresion

For the ICC dataset (SNP-h^2^=13%), no SNPs reached genome-wide signficance^30^. Instead, we focus on those SNPs reaching suggestive significance (p<1×10^5^, N=152 SNPs). After clumping (using clump_kb=3000 and clump_r2=0.1 to match PGC thresholds), 19 independent SNPs were retained. The logOR and SE for these 19 SNPs were extracted from the PGC-MDD data for use as genetic instruments when testing the causal effect of cannabis use (the exposure) on depression (the outcome).

As both cannabis use and depression are binary outcomes, the effect estimates represent the odds for the outcome per unit increase in the log OR for the exposure. To assist with interpretation, these effect estimates were converted (by multiplying log ORs by 0.693 and then exponentiating) to give the OR per doubling in odds of the binary exposure^50^.

## Results

### Univariate analyses within UK Biobank

Of the 126291 individuals with both mental health questionnaire and genetic data available, 43.8% are male, 60.2% have ever smoked and 60.9% are high frequency alcohol drinkers. Table 1 shows the prevalence of each of the seven phenotypes of interest. For cannabis use phenotypes, cases are more likely than controls to be male and high frequency alcohol users. For depression and self-harm phenotypes, cases are less likely than controls to be male and high frequency alcohol users. For all phenotypes, cases more frequently report smoking (Supplementary Materials Section 1)

**Table 1:** Prevalence of traits in UK Biobank.

| Phenotype | Control | Case | SNP-h<sup>2</sup> | SNP-h<sup>2</sup> SE | LD-score intercept | LD-score intercept SE |
| --- | --- | --- | --- | --- | --- | --- |
| Cannabis Use | 97827 | 28282 | 0.119 | 0.011 | 1.038 | 0.007 |
| Initial Use | 97827 | 19202 | 0.084 | 0.010 | 1.012 | 0.007 |
| Continued Use | 97827 | 9080 | 0.135 | 0.016 | 1.013 | 0.006 |
| Depression | 71134 | 30075 | 0.126 | 0.010 | 1.000 | 0.008 |
| Single Episode | 71134 | 12099 | 0.057 | 0.012 | 0.997 | 0.007 |
| Recurrent | 71134 | 17976 | 0.147 | 0.012 | 1.002 | 0.008 |
| Self Harm | 120408 | 5520 | 0.121 | 0.020 | 1.009 | 0.006 |

We performed GWA analyses for each of the seven traits within UK Biobank (Supplementary Materials Section 1) and then using these statistics estimated the SNP-heritability of each trait, using LD-score regression. These results are reported on the liability scale. For both cannabis use and depression, the more severe phenotype (continued cannabis use and recurrent depression, respectively) have higher SNP-heritability estimates. (Supplementary Materials Section 1 includes estimates for over-or under-sampling scenarios).

### Bivariate analyses

#### Phenotypic relationships within UK Biobank

The prevalence of all depression and self-harm phenotypes is significantly higher amongst cannabis users than non-users (Table 2). Furthermore, the associations are stronger for more severe phenotypes (continued vs initial cannabis use and recurrent vs single episode depression), as illustrated in Figure 2. Cannabis use phenotypes are most strongly associated with self-harm. There is a strong relationship between depression and self-harm (OR=10.13, 95% CI=9.40-10.93). These patterns of association were also generally seen in each of the stratified analyses (Supplementary Materials Section 2 has details and exceptions).

**Table 2:** Phenotypic relationship between traits within UK Biobank.

| Predictor | Outcome | OR | 2.5% | 97.5% | p |
| --- | --- | --- | --- | --- | --- |
| Cannabis Use | Depression | 1.64 | 1.59 | 1.70 | 1.19e-213 |
| Initial Use |  | 1.50 | 1.45 | 1.56 | 2.49e-105 |
| Continued Use |  | 1.98 | 1.88 | 2.08 | 7.04e-161 |
| Cannabis Use | Single episode | 1.40 | 1.33 | 1.46 | 1.29e-47 |
| Initial Use |  | 1.33 | 1.26 | 1.40 | 8.64e-26 |
| Continued Use |  | 1.56 | 1.45 | 1.68 | 1.60e-33 |
| Cannabis Use | Recurrent | 1.82 | 1.76 | 1.89 | 1.30e-224 |
| Initial Use |  | 1.63 | 1.56 | 1.70 | 4.03e-108 |
| Continued Use |  | 2.28 | 2.16 | 2.42 | 2.74e-179 |
| Cannabis Use | Self Harm | 2.85 | 2.69 | 3.01 | 3.46e-304 |
| Initial Use |  | 2.38 | 2.23 | 2.54 | 1.26e-148 |
| Continued Use |  | 3.89 | 3.61 | 4.19 | 3.25e-281 |

**Figure 1:**
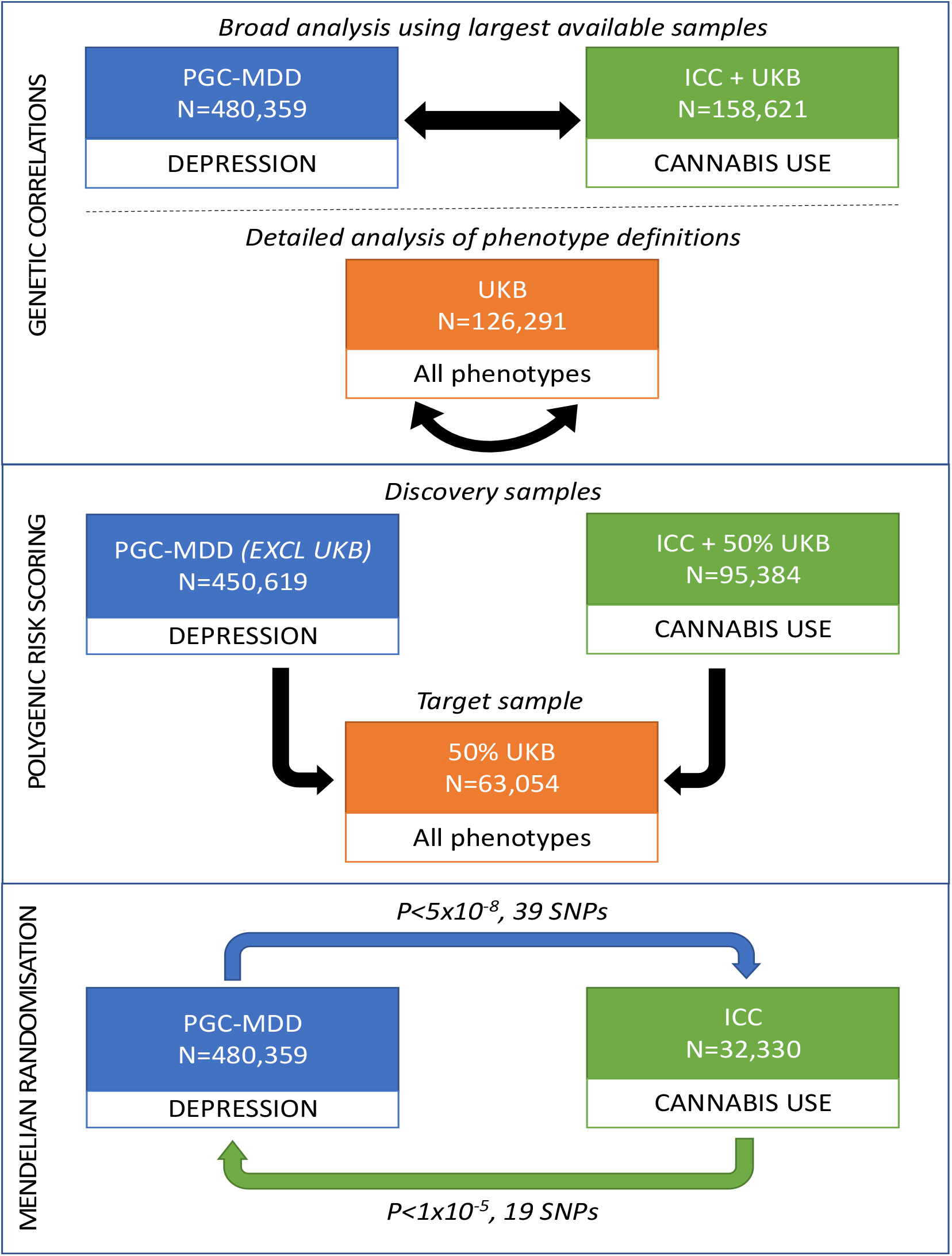
Bivariate Genetic Methods Overview. PGC-MDD = Psychiatric Genomics Consortium Major Depressive Disorder, including 23andMe; ICC = International Cannabis Consortium; UKB = UK Biobank

**Figure 2:**
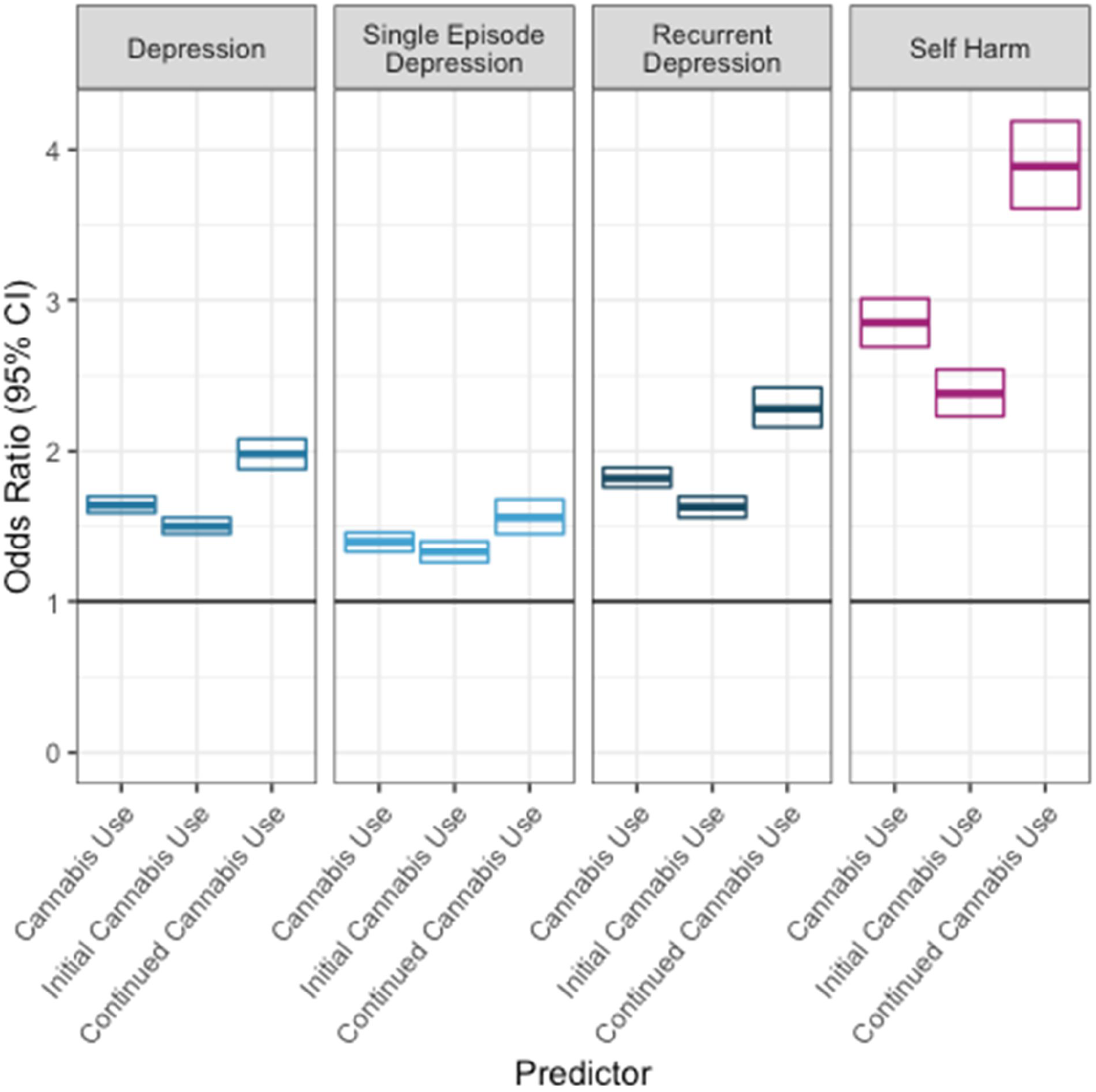
Phenotypic relationship between traits within UK Biobank

The phenotypic findings do not consider the direction of effects; it cannot be determined whether one factor is on the causal pathway to another or if there are upstream factors acting on all of the phenotypes under consideration.

### Genetic correlations

#### Maximum available sample size

We find a genetic correlation between cannabis use and depression of rg=0.289, SE=0.036, p=1.45×10^−15^ using our meta-analysis of ICC+UKB (N=158,621, SNP-h^2^=0.116, SE=0.009) and summary statistics made available by the PGC-MDD and 23andMe (N=480,359, SNP-h^2^=0.087, SE=0.004).

#### Detailed phenotypes within UK Biobank

We observe high within-trait genetic correlations between initial and continued cannabis use (rg=0.885, SE=0.083, p=1.03×10^−26^) and single episode and recurrent depression (rg=0.941, SE=0.098, p=1.02×10^−21^). In cross-trait analyses, as with phenotypic observations, all cannabis use traits are modestly and significantly correlated with depression traits (rg=0.255-0.345, p=1.16×10^−2^-2.36×10^−7^) and self-harm (rg=0.298-0.308, p=1.17×10^−3^-5.44×10^−5^). Depression phenotypes show a high genetic correlation with self-harm (rg=0.681-0.745, p=8.21×10^−7^-1.68×10^−20^). However, in contrast to our phenotypic findings, cross-trait genetic correlations do not show the same pattern of varying magnitude across different definitions of cannabis use and depression within UK Biobank (Figure 3, Supplementary Materials Section 3).

**Figure 3:**
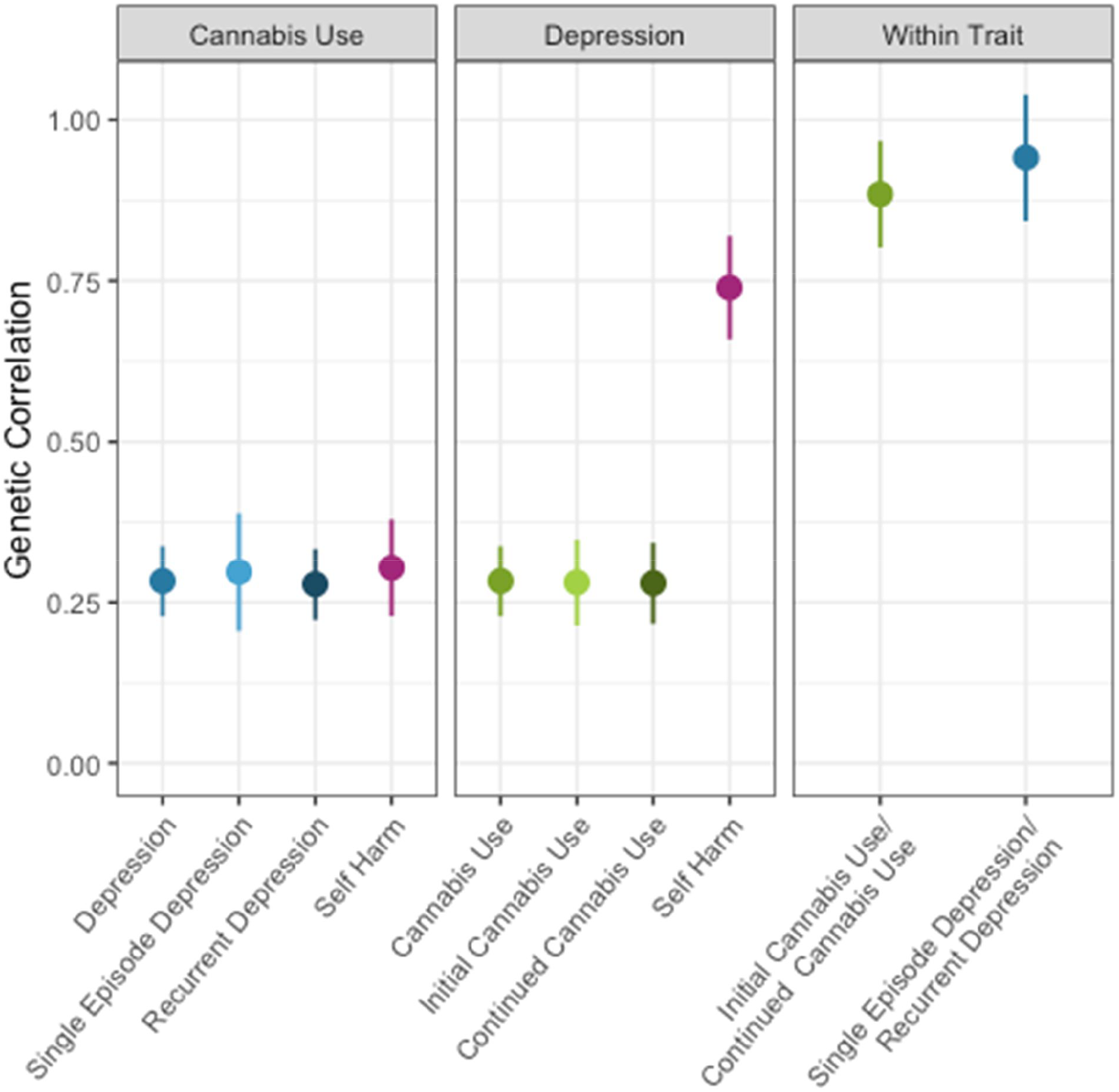
Genetic Correlations within UK Biobank, calculated using LD-score regression

### Polygenic risk scores

The two calculated polygenic risk scores for depression and cannabis use were tested against the seven phenotypes of interest in the 50% UK Biobank target sample (Figure 4). The results report P_threshold_=0.3 as a representative value, full findings in Supplementary Materials Section 4. A Bonferroni correction for the two PRS and 7 traits (which does not consider the correlation between these traits) gives a significance threshold of p=0.05/(2×7), or p=3.57×10^−3^.

**Figure 4:**
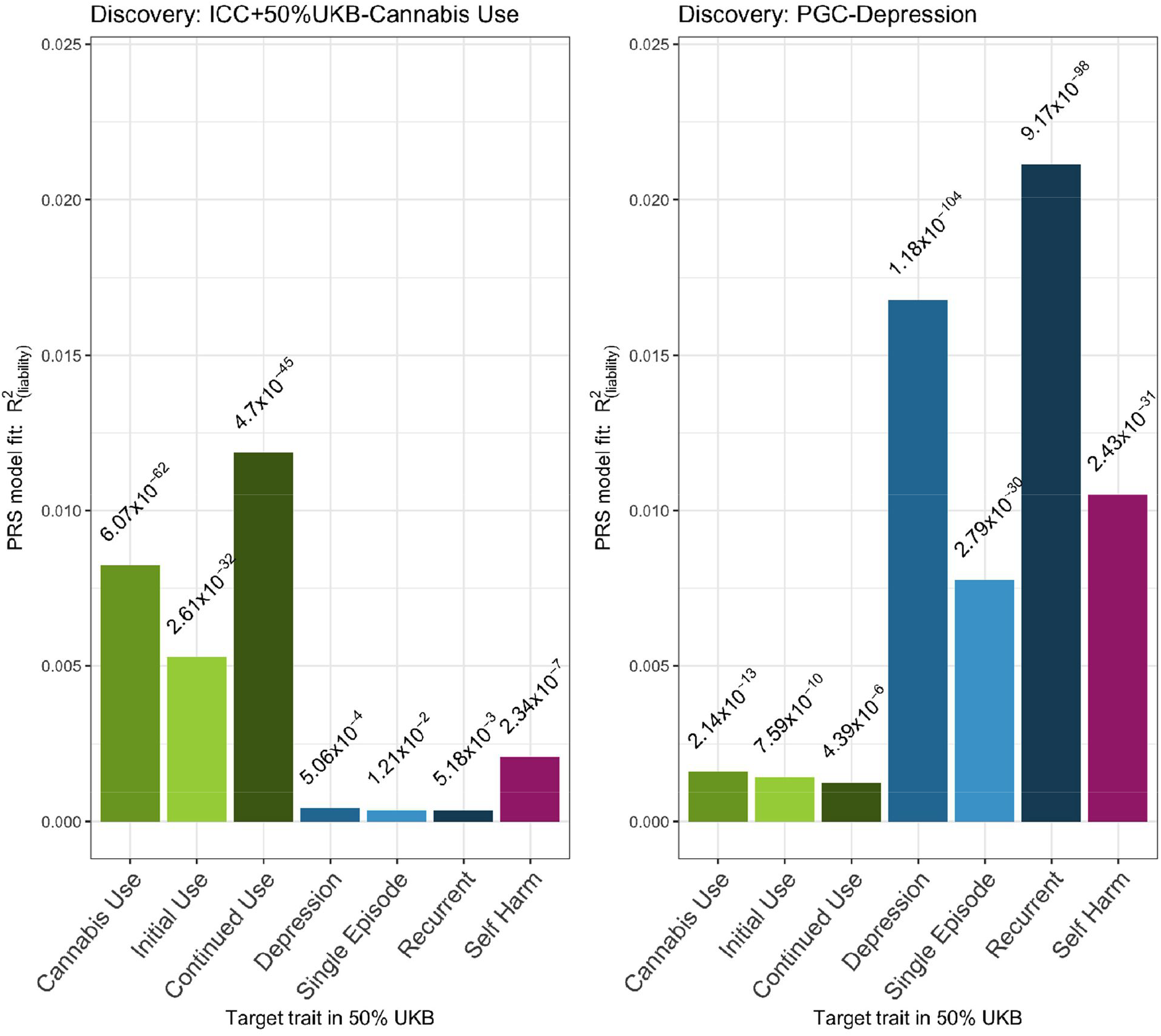
Polygenic risk scoring in UK Biobank

#### Polygenic risk for cannabis

Looking within-trait, PRS_(CANN)_ accounts for a significant proportion of the variance in cannabis use in the UK Biobank target sample (R^2^=0.82%, p=6.071×10^−62^). PRS_(CANN)_ explains a greater proportion of variance in continued cannabis use (R^2^=1.2%, p=4.697×10^−45^) than initial cannabis use (R^2^=0.53%, p=2.609×10^−32^). In cross-trait analyses, PRS_(CANN)_ accounts for a small proportion of variance in depression (R^2^=0.043%, p=5.058×10^−4^) and does not reach significance when considering single episode (R^2^=0.037%, p=1.211×10^−2^, NS) or recurrent depression (R^2^=0.037%, p=5.180×10^−3^, NS). PRS_(CANN)_ predicts a greater proportion of variance for self-harm than depression (R^2^=0.21%, p=2.337×10^−7^).

#### Polygenic risk for depression

PRS_(DEP)_ accounts for a significant proportion of variance in depression in the UK Biobank target sample (R^2^=1.7%, p=1.176×10^−104^) and accounts for more variance in recurrent depression (R^2^=2.1%, p=9.173×10^−98^) than single episode depression (R^2^=0.78%, p=2.792×10^−30^). PRS_(DEP)_ also accounts for R^2^=1.1% of the variance in self-harm (p=2.427×10x^−31^). Looking at cannabis phenotypes, PRS_(DEP)_ accounts for a significant but smaller proportion of the variance in cannabis use (R^2^=0.16%, p=2.144×10^−13^), initial cannabis use (R^2^=0.14%, p=7.589×10^−10^) or continued cannabis use (R^2^=0.13%, p=4.393×10^−6^).

Results are consistent with PRS_(CANN)_ being less well powered than PRS_(DEP)_ due to smaller discovery sample size, but overall, findings broadly reflect those patterns identified using phenotypic and LD-score approaches, showing that there is evidence of association between cannabis and depression phenotypes and each of these traits are also related to self-harm.

### Mendelian randomisation

Two-sample MR analyses results are shown in Table 3, and Supplementary Materials Section 5.

**Table 3:** Two-sample Mendelian randomisation results.

| Exposure-Outcome | Method | Estimate | OR | SE | p |
| --- | --- | --- | --- | --- | --- |
| Depression-Cannabis | IVW | 0.150 | 1.109 | 0.114 | 0.188 |
|  | MR-Egger | 0.283 | 1.217 | 0.720 | 0.694 |
|  | (intercept) | -0.004 | 0.997 | 0.023 | 0.851 |
| Cannabis-Depression | IVW | 0.019 | 1.013 | 0.013 | 0.167 |
|  | MR-Egger | 0.002 | 1.001 | 0.038 | 0.968 |
|  | (intercept) | 0.003 | 1.002 | 0.006 | 0.632 |

#### Causal effect of depression on cannabis use

An MR analysis using 39 SNPs previously association with depression (reaching genome-wide significance in the published PGC-MDD and 23andMe GWA) indicated no evidence of association with cannabis use, using the ICC GWA cannabis use summary statistics. Both IVW and Egger models did not reach significance (p>0.01), with no evidence of directional horizontal pleiotropy, as indicated by the MR-Egger intercept.

#### Causal effect of cannabis use on depression

Nineteen independent SNPs associated with cannabis use (reaching suggestive significance p<1×10^−5^ in the published ICC cannabis use GWA) were used to test for evidence of cannabis use having a causal effect on depression using the PGC-MDD and 23andMe GWA summary statistics. Again, both IVW and Egger models did not reach significance (p>0.01), with no evidence of directional horizontal pleiotropy.

## Discussion

### Summary of findings

In UK Biobank, cannabis use is associated with an increased likelihood of depression and particularly self-harm. There is also a genetic overlap between these traits, as demonstrated using cross-trait LD-score regression. Polygenic risk for depression is associated with depression, self-harm and cannabis use, however cross-trait results using polygenic scores for cannabis use are more equivocal. We explored how these phenotypic and genetic relationships vary for initial or continued cannabis use and for single episode or recurrent depression. Phenotypically, cross-trait associations are stronger for more severe trait subtypes. However, this pattern is not evident in the genetic analyses. Finally, as these analyses cannot determine the direction of effects, Mendelian randomisation was implemented using published GWA summary statistics. We did not find evidence to support causality between cannabis use and depression in either direction, although we note that these MR analyses are underpowered.

### Phenotypic and genetic relationships between cannabis use, depression and self-harm

Phenotypic associations between cannabis use and depression are significant but the association between cannabis use phenotypes and self-harm is stronger than those observed with depression, mirroring reports in other populations^16,19,23^. Stratified analyses suggest that these associations are not driven by differences in sex, smoking or alcohol consumption.

Significant genetic correlations between cannabis use, depression and self-harm are also in line with some^15,23,24^, but not all^16^ previous findings in family studies. The genetic correlation between cannabis use and depression estimated here is higher than those previously published using the same methodology^13^, likely reflecting the increased power of more recent depression GWA analyses. Indeed, the genetic correlation observed here between cannabis use and depression also exceeds that reported between cannabis use and schizophrenia (rg=0.245, SE=0.031, p=5.81×10^−15^)^13^.

Polygenic risk scoring methods show that PRS_(DEP)_ accounts for a small but significant proportion of the variance in each of the cannabis phenotypes. The PRS_(CANN)_ is less powerful (shown by within-trait prediction) and whilst significantly predicting depression, fails to reach significance for either single episode or recurrent depression. Both PRS_(DEP)_ and PRS_(CANN)_ account for a significant proportion of the variation in self-harm.

Significant PRS show the importance of cross-trait polygenic effects, however they explain a very small proportion of trait variance. This is expected; PRS is an out-of-sample prediction method and current GWA samples have limited power to capture the small genetic effects for complex traits like cannabis use and depression. As GWA sample sizes continue to grow, we predict polygenic risk scores should explain an increasing proportion of trait variance, and so will become more useful as a tool for individual prediction. In the meantime, the convergent genetic results from both LD-score regression and polygenic risk scoring methods both indicate shared genetic influences on cannabis use, depression and self-harm that warrant further attention.

### The impact of trait definitions

Whilst all phenotypic relationships tested are significant, our observation that associations are stronger for more severe trait definitions is consistent with previous reports^16,19^. Our SNP-h^2^ estimates of depression and cannabis use are also inline with previous findings^13,14^, but we show that continued cannabis use is a more heritable trait than initial cannabis use and that there is a strong genetic correlation between these two. This suggests that focussing on continued cannabis use may be beneficial for genetic analyses.

Unlike phenotypic patterns, the strength of genetic relationship does not vary with the trait definition, using LD-score or PRS cross-trait approaches. This may reflect a difference in the phenotypic versus the genetic relationships for these traits, however it may also be due to limitations in the statistical power of genetic analyses to differentiate between subtypes.

Irrespective, there is still substantial evidence of both phenotypic and genetic associations across all traits tested; even initial cannabis use is associated with an increased risk of depression and self-harm, underscoring the importance of work examining the causal mechanisms linking these phenotypes.

### Directions of causality between cannabis use and depression

The direction of causality is of particular interest, given current debates surrounding the legalisation and safety of using cannabis. Longitudinal approaches have been inconclusive^5,9^ and MR offers an alternative genetic-based method to test causality. Here we found no evidence for causality in either direction, but our analyses were underpowered, relying on suggestive but not significant SNPs as instrumental variables for cannabis use. This is in part because we were unable to include the genome-wide significant variants implicated in most recent and largest cannabis GWAS^13^, given issues of bias and sample overlap.

Determining the direction of causality linking cannabis use and depression is key to understanding the potential risks of cannabis use, with associated health policy implications. It is most likely that there is a network of both genetic and environmental factors linking cannabis use, depression and self-harm which will vary between individuals.

## Conclusions

Here we show that cannabis use is phenotypically and genetically associated with depression and self-harm. Whilst the phenotypic relationship is stronger for more severe phenotypes, there is a significant increase in risk for those who use cannabis less than ten times. However, we are unable to draw conclusions regarding the direction of causality linking cannabis use and depression. As has previously been noted, there is more commonly a focus on the relationship between cannabis and psychosis-related phenotypes^9^. This work highlights that when investigating the associations of cannabis use with mental health, both depression and self-harm are important and prevalent traits to consider.

## Acknowledgements

The study represents independent research funded by the National Institute for Health Research (NIHR) Biomedical Research Centre at South London and Maudsley NHS Foundation Trust and King’s College London. The views expressed are those of the author(s) and not necessarily those of the NHS, the NIHR or the Department of Health and Social Care. High performance computing facilities were funded with capital equipment grants from the GSTT Charity (TR130505) and Maudsley Charity (980). Analysis using UK Biobank data was conducted under UK Biobank application number 18177. We would like to thank the research participants and employees of 23andMe for making this work possible. We also thank the members of the International Cannabis Consortium for making summary data available, enabling this work.

